# Effects of prediction and attention on tactile precision in somatosensory gating

**DOI:** 10.64898/2026.03.05.709823

**Authors:** Pierangelo N. D’Onofrio Pacheco, Eckart Zimmermann

## Abstract

Tactile sensitivity is reduced when the limb is in motion, a phenomenon known as somatosensory gating. In a previous study, we demonstrated that discrimination precision but not perceived intensity differed between active and passive movements. Here, we asked whether and how spatial attention modulates tactile precision in active and passive movements. Participants judged the relative intensity of two vibrations while the arm was still, actively moved, or passively transported by a movable platform. Visual attention was directed either to the movement start or goal position. Perceptual bias was reduced during both active and passive movement, independent of attentional allocation. In contrast, precision remained stable during active movement but declined during passive movement when attention was directed to the movement start. However, when attention was focused on the movement goals, precision was also high when doing passive movements. These findings indicate that during active movements, predictions based, likely on an efference copy, ensure tactile precision, whereas passive movements require spatial attention directed to the movement goal.

## Introduction

Tactile sensitivity declines when the limb is in motion, a phenomenon commonly referred to as somatosensory gating. This reduction in perceived intensity has been attributed to both central mechanisms, such as motor preparation and command signals, and peripheral factors such as reafferent masking(Angel & Malenka, 1982; Chapman et al., 1987; Williams & Chapman, 2002). Early electrophysiological studies reported reduced amplitudes of somatosensory-evoked potentials (SEPs) around movement onset, suggesting that gating operates at both spinal and cortical levels (Seki & Fetz, 2012; Starr & Cohen, 1985).

In our previous work (D’Onofrio Pacheco PN, 2026)**(D’Onofrio Pacheco & Zimmermann, 2026)**, we demonstrated that reductions in perceived intensity during active movement are statistically indistinguishable from those observed during passive movement. Crucially, however, tactile discrimination sensitivity differed between these conditions: precision was preserved during active movements but reduced during passive movements. Additional experiments showed that the predictability of movement kinematics can account for this difference, because when participants could anticipate the velocity of passive motion, the brain could generate more accurate predictions about forthcoming sensory input, allowing it to separate task-relevant tactile signals from movement-related noise. These findings indicate that tactile gating comprises separable components and that perceptual precision depends on the availability of predictive information.

A growing body of research further suggests that gating is modulated by contextual and cognitive factors. Manipulations of task relevance, timing, and sensory predictability can reduce or even reverse suppression, enhancing discrimination at behaviorally relevant locations (Colino et al., 2014; Gertz et al., 2017; Kilteni & Ehrsson, 2022; Clare Press et al., 2023; Voudouris et al., 2019). Moreover, studies conducted with stationary limbs show that directing attention to a specific body location enhances tactile processing and modulates somatosensory cortical activity (Gherri et al., 2023; Sambo & Forster, 2011; Schweisfurth et al., 2014). Visual spatial attention can also bias tactile processing at corresponding external locations(Driver & Spence, 1998; Eimer, 2001), and visuo–tactile interactions are strongly influenced by attentional allocation across modalities (Badde et al., 2020; Göschl et al., 2014). Together, these findings suggest that perceptual precision during movement may depend not only on motor-based predictions but also on whether visual attention is directed to a spatial location that provides reliable information about the upcoming movement, thereby enhancing the predictability of its sensory consequences.

In this study, we tested the influence of overt visual attention on intensity estimates of tactile stimuli presented during limb movement. Participants directed gaze either to the start or to the goal location of a horizontal arm movement while tactile stimuli were delivered during active or passive displacement of the arm. In the active condition, detailed information about movement kinematics was available in the internal motor commands. An efference copy of these commands may be used to continuously predict and evaluate reafferent tactile input. In the passive condition, however, no efference copy is available, and the brain must rely on learned externally derived signals to infer movement kinematics. Although both proprioceptive and visual cues can contribute to this inference, vision typically provides a more reliable estimate of the movement endpoint (Ernst & Banks, 2002; Körding & Wolpert, 2004).

We therefore reasoned that directing gaze to the goal location would provide the visual system with direct spatial information about the movement goal. This enhanced spatial information could improve the brain’s predictions about the ongoing passive movement and its sensory consequences. Conversely, directing gaze to the movement start position would draw attentional resources away from the goal and reduce the reliability of such spatial predictions. We hypothesized that perceived intensity would be attenuated in all movement conditions, reflecting a general movement-related bias, whereas discrimination precision would remain high and attention-independent during self-generated movement but decline during passive movement unless attention was focused at the goal location.

## Methods

### Participants

Eighteen right-handed adults (12 females; mean age = 22.2 years, SD = 3.1) participated. All reported normal or corrected-to-normal vision, right handed and no history of neurological or psychiatric disorders. The study conformed to the Declaration of Helsinki and was approved by the ethics committee of the Faculty of Mathematics and Natural Sciences, Heinrich Heine University Düsseldorf.

### Apparatus and materials

Participants sat with the head stabilized by a chin rest 67 cm from a 24-inch monitor (1920 × 1080 px, 144 Hz refresh rate). Eye position was tracked monocularly (right eye) with a Tobii 5 eye tracker (250 Hz), mounted below the screen.

The right arm rested on an ergonomic support. In the active condition, participants moved a Touch X haptic stylus (3D Systems) along a 20 cm horizontal path between two visual targets. In the passive condition, the same stylus was attached to a custom linear rail driven by a stepper motor (Arduino Leonardo controller) that translated the hand at a constant velocity of 200 mm/s.

Tactile stimulation was delivered by a 1 cm vibrotactile actuator fixed over the median-nerve region of the right forearm. Each vibration lasted 300 ms. On every trial, the probe vibration had a fixed duty cycle of 55%, while the comparison vibration was randomly selected from nine possible duty cycles (10, 20, 30, 40, 50, 60, 70, 80, or 90%). Vibrations were triggered by an Arduino Nano microcontroller synchronized with the timing routines. Participants wore Soundcore Life Q30 noise-canceling headphones to mask auditory cues. Responses were given on a three-key keypad (Pimoroni Keybow). All experimental measurements including eye tracking, stylus tracking, stimulus timing, and response collection were controlled by custom C# scripts in a 3D environment.

### Design and procedure

The experiment followed a 3 (Movement: Control, Active, Passive) × 2 (Attention: Start, end) factorial within-subjects design. The order of movement blocks was counterbalanced across participants, and attention conditions were randomized between each block.

In the control condition, participants’ arms remained still throughout the entire condition. After the eye tracker confirmed that gaze was maintained within 1° of the cued fixation location for 200 ms, a probe vibration was delivered, followed 800 ms later by the comparison vibration. In the active movement condition, participants moved the stylus from the start position to the endpoint following the green cue. The probe vibration occurred 200 ms after movement onset, and the comparison vibration was delivered 200 ms after the stylus reached the endpoint, Figure 1 A. In the passive condition, the linear rail transported the participant’s arm along the same trajectory and velocity profile as in the active condition, and the timing of the vibrations was identical Figure 1 B.

**Figure 1:**
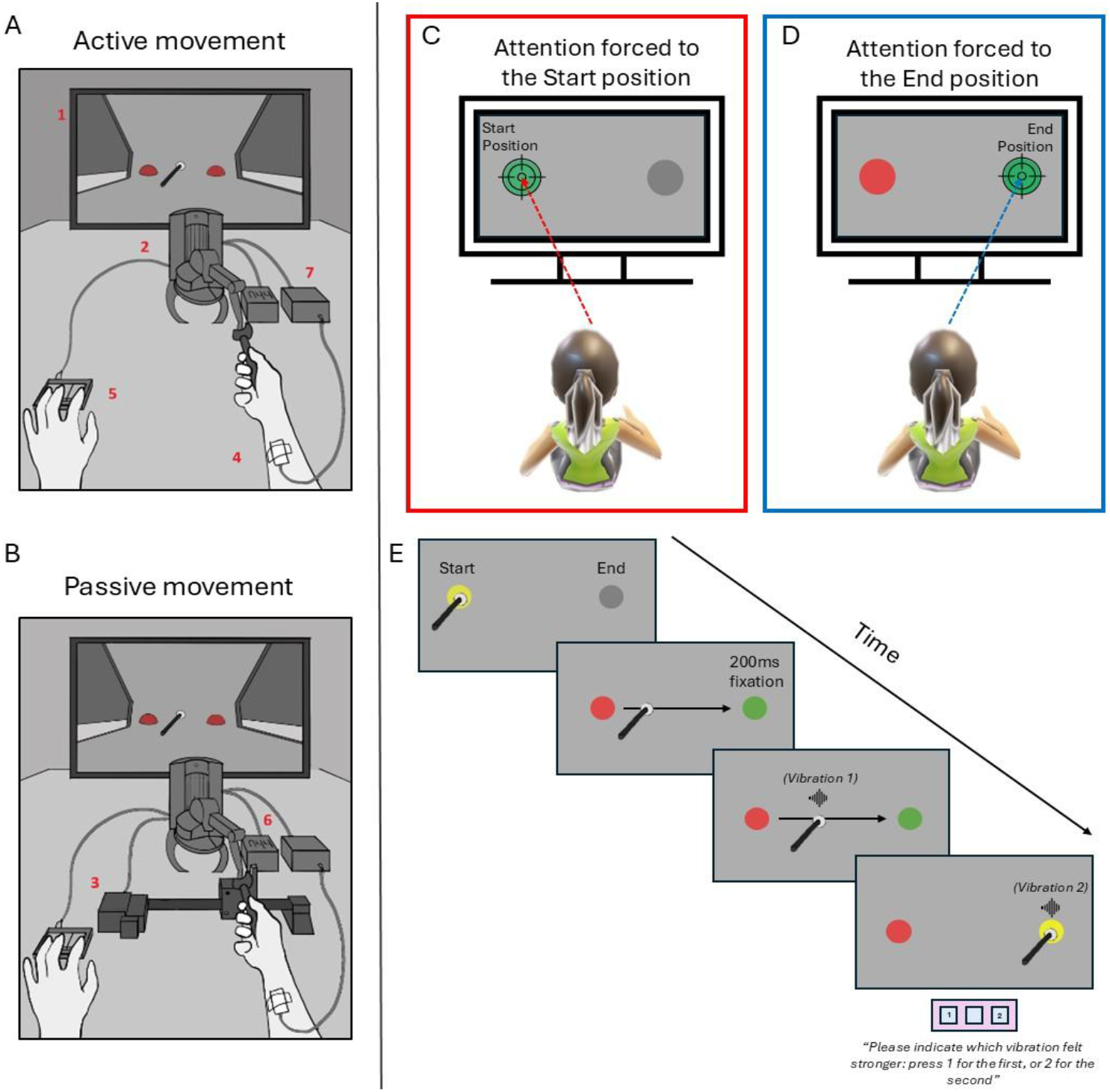
Experimental set-up and task design. (A) Schematic of the active task. Participants moved their right arm with a stylus from the first (left) to the second (right) sphere. A vibrotactile stimulus was delivered on the median nerve with a 200ms delay, same for the passive condition). After reaching the second sphere, a second vibration was presented 200 ms later. Participants judged which vibration felt stronger by pressing a key (“1” = first stronger, “2” = second stronger) on the Pimoroni Keybow(5). (B) Active condition apparatus. The task display (1) guided the movement of a stylus tracked by the Touch X system (2). Vibrotactile stimulation was controlled by an Arduino Nano (7), with the actuator placed over the right median nerve (4). Responses were given via the Keybow. (C) Passive condition apparatus. The participant’s arm was moved by a rail system (3) driven by a stepper motor (Arduino Leonardo, 6). Passive movements (200 mm/s). Vibrotactile stimuli were delivered 200 ms after movement onset and again 200 ms after reaching the target sphere.

At the start of each trial, two colored spheres appeared on the screen corresponding to the start and endpoint of the horizontal movement path. One sphere changed color to indicate the location participants should fixate. The trial began once gaze was maintained within 1° of the cued location for 200 ms, at which point the fixation cue turned green to signal that the trial had started, see figure 1,E. In the attention manipulation, the fixation cue appeared either over the starting sphere (Start condition, Figure 1 C) or the endpoint sphere (Shifted condition, Figure 1 D).

On every trial, participants judged which of the two vibrations felt stronger by pressing “1” if the first vibration felt stronger or “2” if the second did. Each participant completed one block for each movement type, with 100 trials per block, resulting in a total of 500 trials including practice. The eye tracker was recalibrated between blocks whenever drift exceeded 0.5°.

### Data analysis

Responses were fitted with cumulative Gaussian psychometric functions using maximum-likelihood estimation for each participant and condition. The point of subjective equality (PSE) was used to indicate perceptual bias. The just-noticeable difference (JND) was given by the standard deviation of the fitted function, indexing discrimination precision. Trials with movement latency, duration, or path curvature exceeding ±2 SD from a participant’s mean were excluded (< 5% of trials).

PSE and JND values were analyzed with separate repeated-measures ANOVAs (Movement × Attention). Sphericity violations were corrected with Greenhouse–Geisser adjustments. Post-hoc tests used Holm-Bonferroni correction. Effect sizes are reported as partial η^2^ for ANOVAs. Analyses were performed in JASP 0.18.1.0, with α = 0.05.

## Results

### Bias (PSE)

A 3 (Movement: Control, Active, Passive) × 2 (Attention: Start, Shifted) repeated-measures ANOVA (figure 2, left) revealed a main effect of Movement, *F*(2, 34) = 18.30, *p* < .001, η^2^_p_ = 0.52, indicating reduced perceived intensity during movement. Neither the main effect of Attention, *F*(1, 17) = 0.77, *p* = .394, η^2^_p_ = 0.04, nor the Movement × Attention interaction, *F*(2, 34) = 1.59, *p* = .218, η^2^_p_ = 0.09, reached significance.

**Figure 2:**
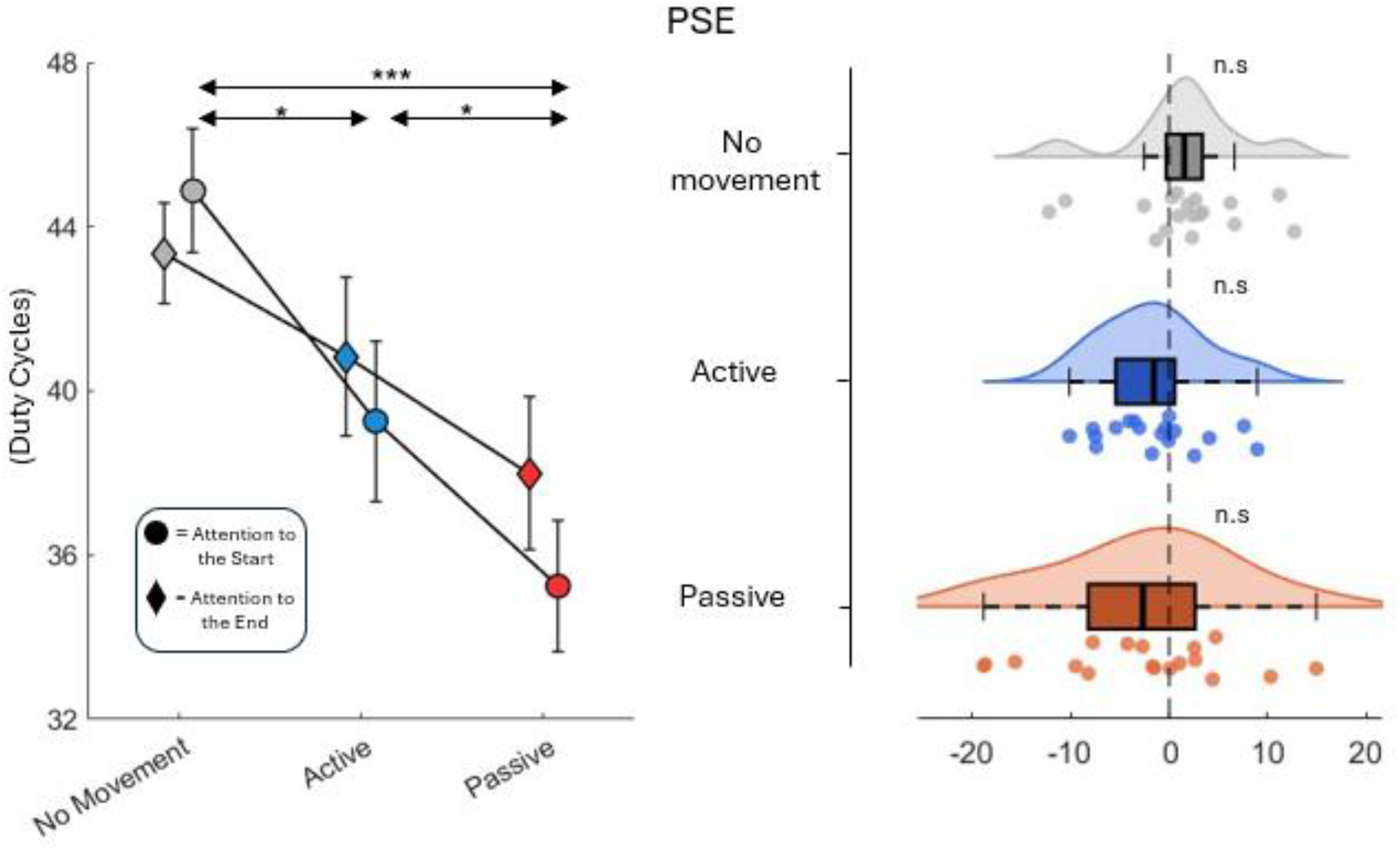
Effects of movement and attention on perceptual bias (PSE). **Left**: Mean PSE values (duty cycles) plotted as a function of Movement condition (No movement, Active, Passive) and Attention. Circles indicate attention directed to the Start position, and diamonds indicate attention directed to the End (goal) position. Error bars represent ±1 SEM. Horizontal brackets denote significant post-hoc comparisons from the 3 (Movement) × 2 (Attention) repeated-measures ANOVA. **Right:** Attentional effect on PSE expressed as the difference between End and Start fixation (Δ = End − Start) for each movement condition. Half-Violin plots show the distribution of individual values; boxplots represent median and interquartile range; dots indicate individual participants. Asterisks denote significant post-hoc comparisons (* p < .05, *** p < .001) and “n.s.” denotes non-significant comparisons.

Post-hoc comparisons (Holm-corrected) showed that both movement conditions yielded significantly lower PSEs than the stationary control condition. The Active condition (*M* = 3.96, *SD* = 0.82) and the Passive condition (*M* = 3.65, *SD* = 0.72) both produced smaller PSEs than Control (*M* = 4.41, *SD* = 0.58), *t*(17) = 2.85, *p* = .022, and *t*(17) = 7.56, *p* < .001, respectively. PSEs were also slightly lower in Passive than in Active movement, *t*(17) = 2.72, *p* = .022.

There was no effect of overt attention. Collapsed across movement, mean PSEs were comparable for Start fixation (*M* = 3.98, *SD* = 0.70) and Shifted fixation (*M* = 3.90, *SD* = 0.71), *t*(17) = 0.88, *p* = .394.

### Precision (JND)

The same 3 (Movement: Control, Active, Passive) × 2 (Attention: Start, Shifted) ANOVA on JNDs (fig 3, left) revealed a main effect of Attention, *F*(1, 15) = 17.48, *p* < .001, η^2^_p_ = 0.54, with higher precision when gaze was shifted to the movement endpoint than when it was centered at the start position. Importantly, a significant Movement × Attention interaction was observed, *F*(2, 30) = 6.53, *p* = .004, η^2^_p_ = 0.30, indicating that the attentional benefit differed across movement types.

**Figure 3:**
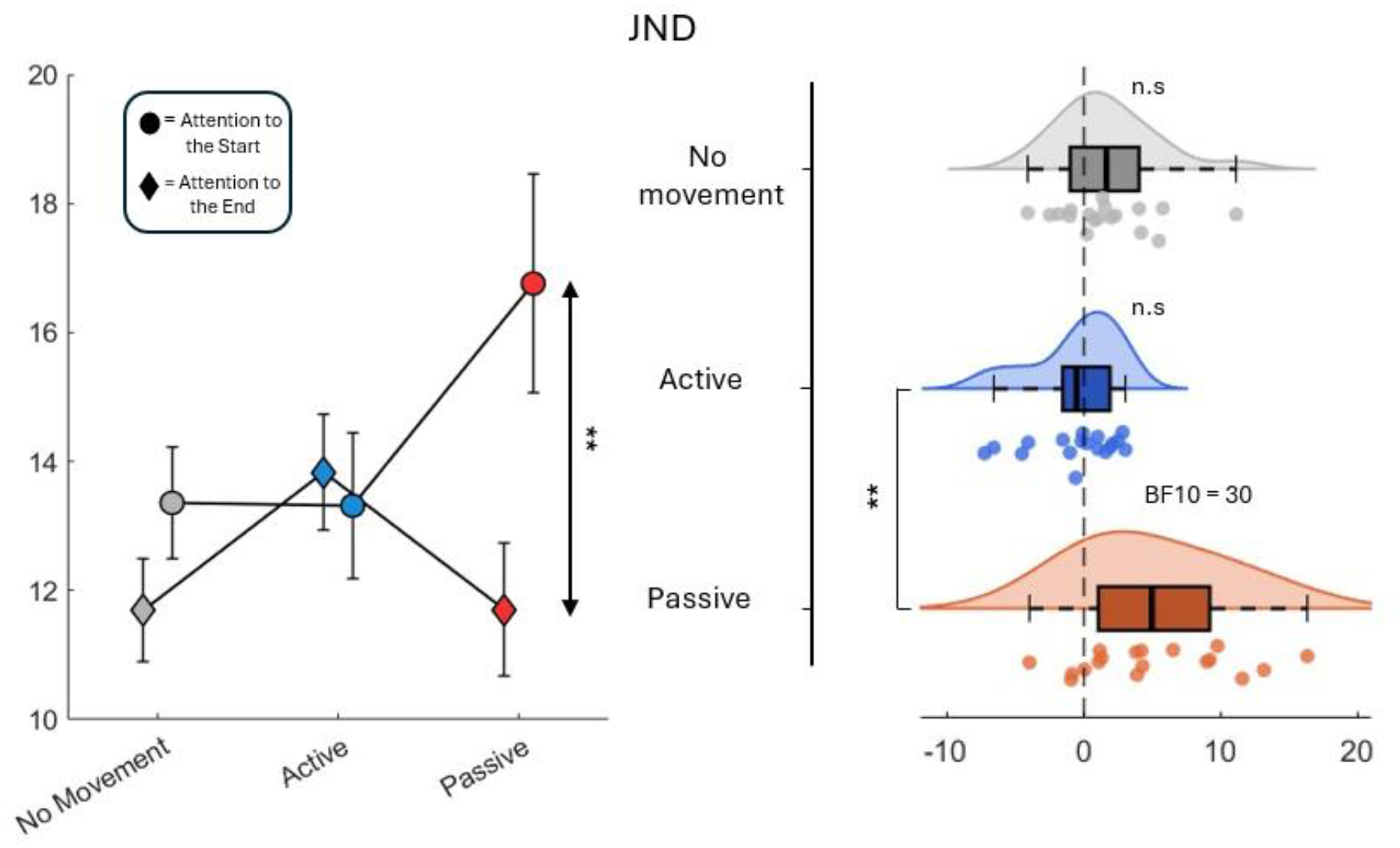
Effects of movement and attention on discrimination precision (JND) **Left:** Mean JND values (duty cycles) plotted as a function of Movement condition (No movement, Active, Passive) and Attention. Circles indicate attention directed to the Start position, and diamonds indicate attention directed to the End (goal) position. Error bars represent ±1 SEM. Vertical brackets denote significant comparisons from the 3 (Movement) × 2 (Attention) repeated-measures ANOVA. **Right:** Attentional effect on JND expressed as the difference between End and Start fixation (Δ = End − Start) for each movement condition. Violin plots depict the distribution of individual values; boxplots show median and interquartile range; dots represent individual participants. Bayesian factor (BF_10_) is indicated where reported.

Post-hoc tests showed that during Active movement, JNDs were low and unaffected by attention (Start: *M* = 1.33, *SD* = 0.48; Shifted: *M* = 1.38, *SD* = 0.38; *p* = 1.00). During Passive movement, however, tactile precision depended strongly on gaze position (Start: *M* = 1.67, *SD* = 0.72; Shifted: *M* = 1.17, *SD* = 0.44; *p* = .019, BF_10_ = 29.22). Notably, the Passive-Shifted condition reached the same precision level as the Active conditions, indicating full recovery of discrimination when attention was directed to the goal.

In contrast, Control trials at rest showed no significant attentional modulation (Start: *M* = 1.34, *SD* = 0.37; Shifted: *M* = 1.17, *SD* = 0.34; *p* = .692).

### Direct quantification of attentional modulation

A one-way repeated-measures ANOVA on the attentional difference scores (Δ = End − Start) for JNDs confirmed a significant effect of Movement condition (F(2, 34) = 6.65, p = .004). Bonferroni-corrected post-hoc comparisons revealed that the attentional benefit was significantly larger in the Passive than in the Active condition (p = .010; Figure 3, right). A Bayesian paired t-test further confirmed that attention improved precision within the Passive condition (BF_10_ = 29.11), providing strong evidence for an attentional effect on discrimination when movement was externally generated. No comparable attentional effect was found in the Active or Control conditions, and attentional manipulation did not influence PSE values in any condition (see Figure 2, right).

## Discussion

The present findings address the mechanisms underlying somatosensory gating by demonstrating that movement differentially affects two aspects of tactile perception: perceptual bias and discrimination precision are differently modulated during limb motion. Consistent with our hypothesis, perceived intensity was attenuated during both active and passive movement, indicating that bias reflects a general movement-related modulation rather than a process specific to motor commands. In contrast, discrimination precision depended critically on the availability of predictive information. Precision remained stable during active movement regardless of attentional allocation, whereas during passive movement it declined unless attention was directed to the movement endpoint.

The different modulation of accuracy and precision directly supports the view that tactile gating is not a unitary suppression mechanism but the outcome of separable processes. The reduction in perceived intensity likely reflects both central gating signals linked to motor commands and peripheral factors such as sensory masking from movement-related afferent inflow (Angel & Malenka, 1982; Chapman et al., 1987; Williams & Chapman, 2002). Crucially, however, precision was preserved only when predictive information about movement kinematics was reliable. During active movement, internal motor commands generate an efference copy that supports forward prediction of reafferent tactile input (Wolpert & Flanagan, 2001), thereby stabilizing discrimination performance. During passive movement, in the absence of such motor-based prediction, precision became dependent on externally derived signals. This interpretation is consistent with recent demonstrations that central predictive mechanisms modulate tactile suppression beyond proprioceptive consequences (Altermatt et al., 2023; Arikan et al., 2024).

Our manipulation of gaze position demonstrates that directing visual attention to the spatial location toward which the limb is moving can compensate for the absence of an efference copy. Directing gaze to the movement endpoint likely increased the reliability of spatial predictions derived from visual input, restoring precision to the level observed during voluntary action. In contrast, maintaining attention at the movement start position reduced the availability of predictive information about the goal, leading to diminished discrimination sensitivity. These findings extend previous work showing that tactile processing is enhanced at attended locations and that visual attention can modulate tactile gain at spatially corresponding positions (Driver & Spence, 1998; Eimer, 2001; Sambo & Forster, 2011). Together with the preserved precision during active movement, these results point to two routes for maintaining tactile fidelity during motion: an automatic, agency-based route in which efference copy stabilizes sensory processing, and a compensatory route in which visual attention to the movement goal supplies the spatial predictions that are otherwise absent.

These findings have broader implications for understanding predictive control and agency. Within frameworks that link motor prediction to sensory processing (Friston, 2010; C. Press et al., 2023), our results suggest that the nervous system deploys at least two mechanisms to maintain tactile discrimination during movement: efference-based prediction during voluntary movement, and attention-dependent compensation consistent with goal-directed attentional control (Corbetta & Shulman, 2002) when agency is absent. Conditions characterized by impaired efference copy or altered attentional allocation may disrupt this balance, leading to degraded sensory discrimination (Brown et al., 2013). More generally, the present results demonstrate that tactile gating is not a uniform suppressive process: motor-based prediction preserves discrimination automatically, whereas attention restores it when prediction must be externally sustained.

## Author Contributions

All authors contributed to the study concept and to the design. Stimuli were designed by P.D.P and performed the data analysis. All authors contributed to the interpretation of results. P.D.P. drafted the manuscript, and E.Z. provided critical revisions. All authors approved the final version of the manuscript for submission.

